# Modeling viral shedding and symptom outcomes in oseltamivir-treated experimental influenza infection

**DOI:** 10.1101/2025.05.02.651873

**Authors:** Yasuhisa Fujita, Marwa Akao, Daiki Tatematsu, Shingo Iwami, Naotoshi Nakamura, Shoya Iwanami

## Abstract

Influenza remains a global public health concern, and although the antiviral drug oseltamivir is widely used to treat infections, questions regarding its actual antiviral efficacy and clinical benefits remain. Here, we evaluated the effects of oseltamivir on viral shedding dynamics in the context of experimental influenza infection. We analyzed individual participant data, including viral load, time to symptom alleviation, and laboratory test measurements, obtained from three publicly available clinical trials involving experimental infections with influenza A and B viruses. We applied mathematical modeling and estimated parameters using a nonlinear mixed-effects model to capture viral infection dynamics. Our analysis revealed that, compared with placebo groups, the oseltamivir-treated groups tended to have lower values in terms of viral load area under the curve, duration of infection, peak viral titer, and time to peak; however, most of these differences were not significant; and no dose-dependent effects were observed. Moreover, there was no significant correlation between time to symptom alleviation and viral load. Some laboratory test parameters showed opposing correlations with symptom-related and viral load-related outcomes. These findings are consistent with distinct mechanisms underlying the symptom-alleviating effects of oseltamivir and its antiviral activity. Our findings suggest that the availability of individual-level data for public use is essential because it enables the evaluation of mechanisms in clinical trials and the development of more appropriate outcome measures.

## Introduction

Influenza is a primary global health concern, causing three to five million cases of severe illness and 290,000–650,000 deaths annually [1]. Seasonal influenza outbreaks significantly burden healthcare systems, particularly affecting high-risk populations, such as older adults, young children, pregnant women, and immunocompromised individuals [2, 3]. While annual vaccination remains the primary preventive strategy, antiviral treatments are essential adjunctive therapies, particularly for severe infections or vaccine mismatches with circulating strains.

Oseltamivir (Tamiflu), a neuraminidase inhibitor (NAI), was first approved in 1999 and has since become one of the most widely prescribed antivirals for influenza treatment and prophylaxis [4]. Despite its widespread use as a medical countermeasure against influenza infection, questions regarding its real-world antiviral efficacy and clinical benefits remain unresolved [5–11]. Because of concerns about its efficacy, oseltamivir was transferred from the World Health Organization’s core list of essential medicines to the complementary list in 2017, and its maintenance on the list is under discussion [12–14]. The United States Centers for Disease Control and Prevention issued Emergency Use Instructions (EUI) for oseltamivir against pandemic influenza A viruses and novel influenza A viruses with pandemic potential in July 2024 [15]. While the epidemiological positioning of oseltamivir remains important, consistent evaluation of its clinical efficacy is desired.

Oseltamivir selectively inhibits the influenza neuraminidase (NA) enzyme, which is essential for releasing newly formed virions from infected cells. By blocking viral egress, the drug limits viral dissemination within the respiratory tract, thereby reducing the overall viral burden [16]. Studies in vitro have demonstrated that oseltamivir is a potent inhibitor of influenza A and B NAs, with half-maximal inhibitory concentration (IC₅₀) values between 0.62–1.07 and 10.01–33.11 nM, depending on the strain [17]. Animal models of influenza infection, including murine, ferret, and nonhuman primate models, support these findings of reduced lung viral loads, inflammatory responses, and mortality rates, at least in part [18–33].

Pharmacokinetic studies indicate that the active metabolite oseltamivir carboxylate achieves therapeutic concentrations in human plasma and respiratory tissues, confirming its systemic antiviral activity [34].

The primary endpoint in pivotal oseltamivir clinical trials was time to alleviate symptoms, which assessed the ability of oseltamivir to shorten the duration of illness [35, 36]. This duration was defined as the interval from treatment initiation until all influenza-related symptoms (fever, headache, muscle aches, cough, sore throat, nasal congestion) were alleviated for at least 24 consecutive hours without additional symptomatic medication. Meta-analyses revealed a 25.2-h reduction in time to alleviate symptoms in adult populations and a 17.6-h decrease in the duration of illness in pediatric populations [7, 8]. However, some studies may not have observed these differences, and thus caution should be exercised in their interpretation [37]. In particular, treatment with oseltamivir and virologic endpoints is also a concern because of variability due to measurement and other factors, and its therapeutic efficacy as an antiviral agent is under debate [38]. Hooker and Ganusov reanalyzed changes in viral load in clinical trials of treatment with oseltamivir for three experimental infections published by a pharmaceutical company [39]. Comparisons of models predicting the start or end of viral shedding suggested an impact of oseltamivir treatment on abnormal viral shedding. The need for data publication was also asserted by comparing the results of clinical trials reported as peer-reviewed papers to confirm consistency.

The present study aimed to evaluate oseltamivir therapy as an antiviral drug using viral shedding dynamics based on a mathematical model of viral infection. Viral shedding kinetics and symptom relief were compared to investigate the relationship between these key endpoints. In addition, we investigated the relationship between available individual laboratory data and endpoints in clinical trials and openly shared the organized data. Through our analysis of three experimental infection trials, we provide one direction for the evaluation of antiviral treatment of influenza.

## Results

### Digitization of published data from oseltamivir clinical trials

To determine the efficacy of oseltamivir in experimental influenza infection trials [40, 41], data from various tests were extracted from published clinical trial reports [6]. Laboratory data, body temperature, time to alleviation of symptoms, age, and sex of 257 participants were obtained from three clinical trials to evaluate treatment efficacy for participants infected with human influenza virus A/Texas/91 (H1N1) (Flu A study) and human influenza B/Yamagata/16/88 virus (Flu B studies A and B) (**Materials and Methods** and **Tables S1– S7**). Commonly measured items were selected and aggregated to ensure consistent analysis for the Flu A and Flu B studies. We selected 24 items from laboratory tests comprising hematologic and biochemical data and treated them as data measured before and after virus inoculation (**Table S7** and **Figure S1**). Owing to differences in protocol, body temperature on day 8 after virus inoculation was not measured in the Flu A study. The items used in the subsequent analysis, excluding viral load, are summarized in **Table S7**. The dynamic change in the viral load for individual participants was obtained from previously published data [39]. Data from 154 of the 257 participants whose viral load values were less than 3 points above the detection limit were excluded when the parameters for the mathematical model described in the next section were estimated (**Figure 1A**). The data from 63 participants who were excluded from the efficacy analysis in the clinical trial with experimental influenza infection were also included in the 154 participants excluded here. For 101 participants, excluding two participants with missing laboratory test data, we compared the distributions of their background information and intervention methods across trials. Flu B study A and Flu B study B were treated without distinction for the following analysis because the observations were made with almost identical protocols. The proportion of female participants was lower in the Flu A study (left panel in **Figure 1B**). Flu A comprised participants less than 35 years old; by contrast, Flu B included participants up to 65 years old, but the mean age was not substantially different between the Flu A and Flu B studies at 22.0 and 29.0 years, respectively (center panel in **Figure 1B**). Dosing regimens differed between the protocols in the Flu A and Flu B studies, but all the implemented dosing regimens remained (right panel in **Figure 1B**). A comparison of laboratory test measurements and body temperatures between the Flu A and Flu B studies revealed that the distributions of several items differed (**Figure 1C**). These differences may be attributed to differences in the measurement methods used and the pathogenicity of the influenza viruses. The differences between trials should be considered when discussing the effects of oseltamivir on these data.

**Fig 1.**
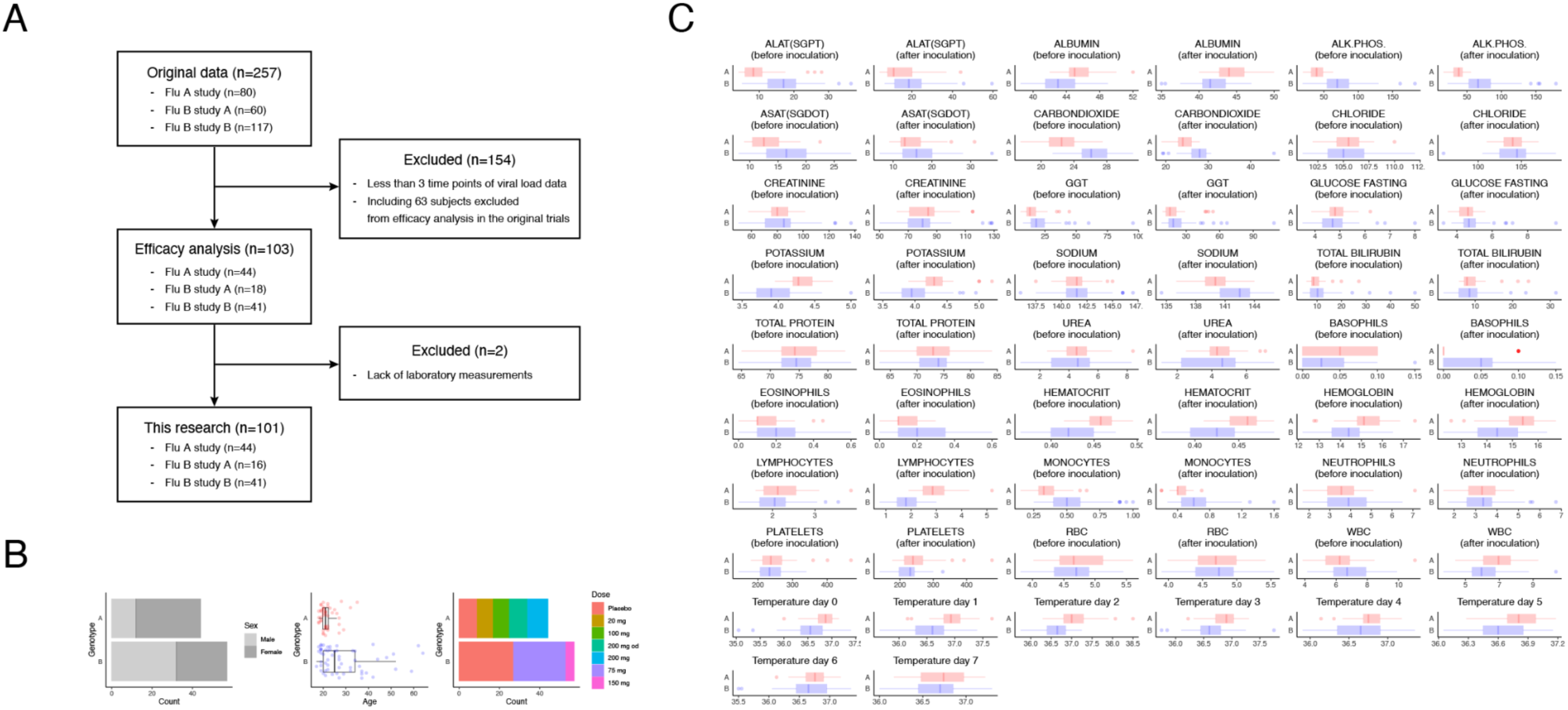
Summary of data obtained from oseltamivir clinical trials. **(A)** Flowchart of participant selection per analysis. Data from participants who met the criteria among data used for efficacy analysis in the human challenge were used for analysis of viral load and correlation analysis using laboratory data. **(B)** Distribution of sex (left), age (center), and dosing (right) by genotype in the analyzed population. **(C)** Summary of laboratory measurements and temperature values by genotype.

### Modeling and quantifying shedding of influenza virus in human volunteers

In clinical trials testing the efficacy of oseltamivir treatment, healthy volunteers were inoculated with a defined amount of influenza A or B virus [40, 41]. In both studies, viral load was measured daily from virus inoculation to evaluate the area under the curve (AUC) of viral load as the primary endpoint. Understanding infection dynamics provides temporal information to assess the antiviral activity of oseltamivir against the growth of influenza viruses in the body. Data from 63 participants were excluded from the efficacy analysis in the Flu A and Flu B studies because the participants did not meet the infection criteria or were suspected of having an infection before the study started (**Table S7**). As described in the section above, 154 participants had detectable virus titers at no more than two time points (**Table S7 and Figure 1A**).

To capture the infection dynamics of the virus, we employed the mathematical model described in the **Materials and Methods** section. We assigned suitable parameters using the nonlinear mixed-effects model (**Figure S2**, **Tables S8 and S9**). Data from participants with fewer than 3 points of detectable viruses were excluded from this analysis of the dynamics of viral shedding. The influenza virus titer varied according to the genotype (**Figure 2A**). Influenza B viruses tended to have a greater AUC (*p* = 1.34 × 10^−3^ by *t* test), longer duration of infection (*p* = 2.33 × 10^−3^ by *t* test), lower peak (*p* = 4.62 × 10^−7^ by *t* test) and later peak time (*p* = 4.76 × 10^−4^ by *t* test) than the influenza A viruses among the analyzed population (**Figure S3**). This difference in viral shedding can be explained by the reduced maximum rate constant for viral replication and death rate of infected cells (*p* = 2.47 × 10^−23^ and *p* = 2.98 × 10^−3^ by Wald test, respectively, **Table S8**) and increased rate constant for infection (*p* = 1.37 × 10^−2^ by Wald test, **Table S8**) of influenza B virus compared with influenza A virus. In previous studies, the AUC and duration of infection were calculated and compared based on measurement values [39]. We compared the reported AUC and duration of virus shedding from treatment initiation with the values estimated by our model (**Figure 2B**). The AUC tended to be smaller, and the duration of infection tended to be greater than the value calculated using the data because measurement error is considered in the statistical model for the estimation. The overall trend of these values was consistent, indicating that coherence was maintained in the evaluation.

**Fig 2.**
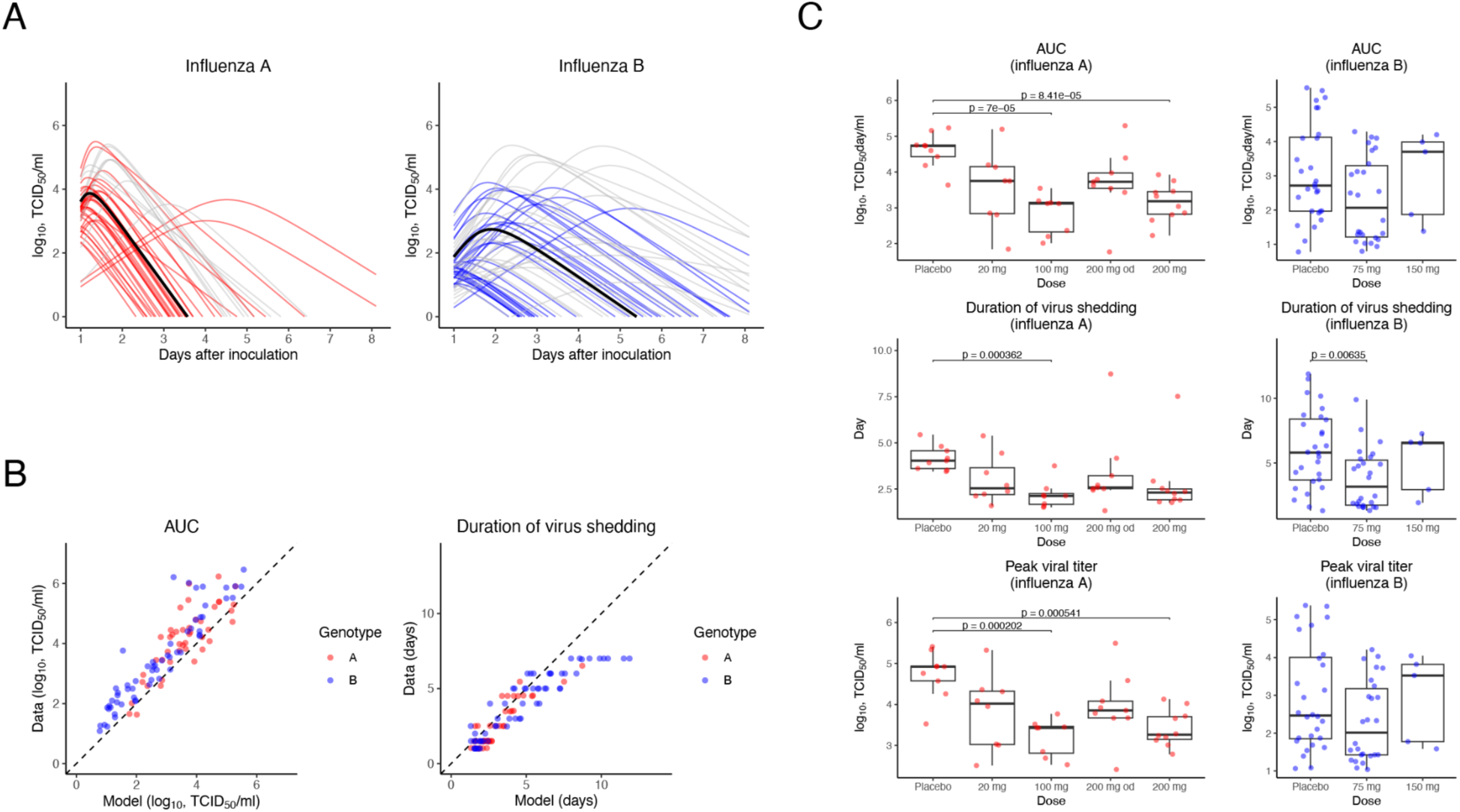
Quantification of the influenza virus A and B loads. **(A)** Dynamics of the viral load after virus inoculation by genotype estimated for individual participants in the oseltamivir-treated (colored) and placebo-treated (gray) groups. The solid and bold black lines represent the population estimates of viral load dynamics. The underlying data and individual fits are shown in **Figure S2**. **(B)** AUC (left) and duration of virus shedding (right) for influenza A (red) and B (blue) infection compared with previous studies by Hooker and Ganusov [39]. **(C)** Comparison of the estimated characteristics of virus dynamics between groups by dose of oseltamivir for influenza A (left) and B (right) viruses. The doses without “od” were administered twice daily; the dose with “od” was administered once daily. We adjusted *p*-values according to the number of group combinations using a post hoc Bonferroni correction.

### Limited effectiveness of oseltamivir treatment on viral load

We investigated the relationship between oseltamivir administration and the characteristics of each participant, estimated by analysis of viral load using a mathematical model of viral infection dynamics. In terms of the summarized features of the changes in viral load, i.e., the AUC, duration of infection, peak viral load, and peak time, there was a trend toward consistent reductions compared with those of the placebo group in each oseltamivir-treated group for both genotypes A and B. Nevertheless, most of the differences were not significant because of the small sample size (**Figure 2C**). More interestingly, no dose-dependent changes in viral load features were observed. A comparison of the estimated parameters of the mathematical model of viral infection dynamics underlying changes in viral load revealed a trend toward an increased rate constant for infection and maximum rate constant for viral replication with oseltamivir administration, but this trend was similarly not dose-dependent (**Figure S4**). Compared with the other model parameters, the estimated death rates of the infected cells showed little change. These results suggested that the infection may have progressed faster, resulting in a reduced viral burden, but the relationship between this progression and oseltamivir administration is unclear.

### There was no significant correlation between viral load and time to alleviation of symptoms

We investigated the relationship between viral load and symptom trends further, which is the basis for the regulatory approval of oseltamivir and viral infection dynamics. Time to alleviate composite symptoms based on the severity of 7 selected symptoms was used as the primary endpoint for treatment of natural infection with oseltamivir [35, 36]. This endpoint was also evaluated in experimental infection studies [40, 41]. We first compared the time to alleviation of symptoms in the placebo and oseltamivir-treated groups for the population we used for our analysis (**Figure 3A**). There was a trend toward a shorter time to alleviation of symptoms in the oseltamivir groups than in the placebo group, as reported, except in the 150 mg b.i.d. group for influenza B virus. However, it should be noted that this is a comparison in a population that excludes data from some of the participants included in the efficacy evaluation in the clinical trial. No significant correlation was detected when the time to alleviate symptoms was compared with the amount of virus per individual calculated using the mathematical model from day 1 to day 8 after virus inoculation (**Figure 3B**). Similarly, no significant correlation with time to alleviation of symptoms was found for the viral load features calculated by the mathematical model (**Figure 3C**). The results of these explorations suggest that the efficacy of oseltamivir in reducing influenza-related symptoms, which is the basis for its approval for influenza treatment and its antiviral effect on viral burden, are due to different mechanisms.

**Fig 3.**
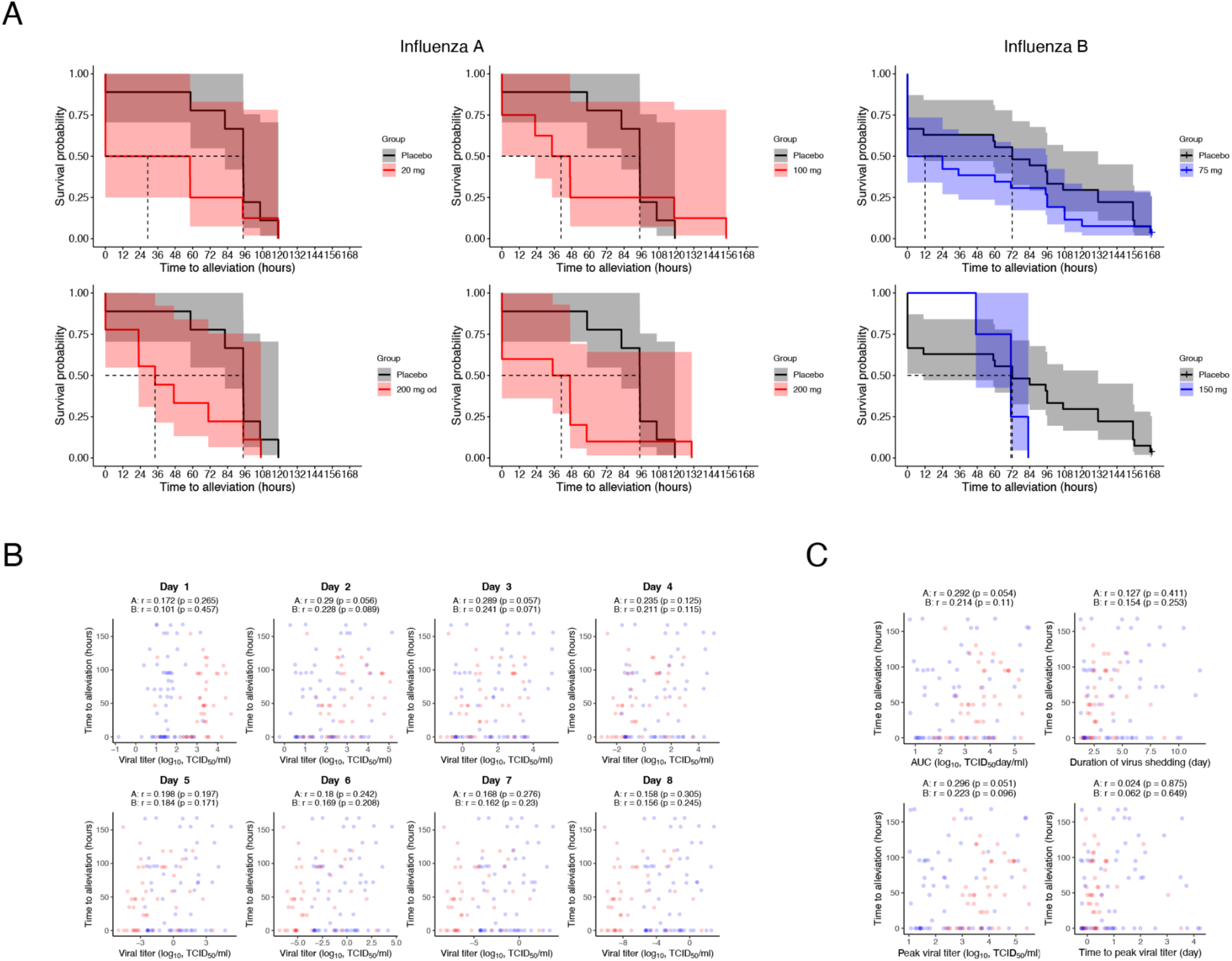
Relationship between time to alleviation and viral infection dynamics. **(A)** Kaplan–Meier survival curves describing the time to alleviation changes for each dose of the oseltamivir treatment group. The doses without “od” were administered twice daily; the dose with “od” was administered once daily. **(B and C)** Correlations between daily viral load **(B)** or viral load-related features **(C)** were calculated using a mathematical model using estimated individual parameters and time to alleviate influenza A (red) and B (blue). Each point represents the value obtained from an individual participant.

### Weak correlations between endpoints and laboratory test measurements suggest contrasting effects of oseltamivir

To determine the effects of oseltamivir on symptoms and viral load, we used laboratory test measurements, including hematologic and biochemical values and body temperature data obtained for safety evaluation in the trials (**Materials and Methods**). As noted in the previous section, there were differences in the distributions of some measured items across studies (**Figure 1C** and **Figure 4A**). No clear correlation was observed between the two primary outcomes, time to alleviate symptoms, the AUC of the viral titer, or any measurements (**Figure 4A**). We calculated partial rank correlation coefficients for each factor to examine its correlation with outcomes, excluding other effects (**Figure 4B**). Notably, some laboratory test items demonstrated opposing correlation directions between outcomes related to symptoms and viral load. For example, chloride levels before inoculation were positively correlated with the time needed to alleviate symptoms. By contrast, the levels were negatively correlated with the AUC, duration of infection, and peak viral titer. Interestingly, the opposite trends were observed after inoculation. Conversely, alkaline phosphatase and carbon dioxide levels were negatively correlated with symptom-related outcomes and positively correlated with viral load-related outcomes before inoculation; however, the correlations were reversed after inoculation. These correlations suggest a potential decoupling between symptomatic relief and viral clearance, indicating that early resolution of symptoms does not necessarily imply a shorter duration of infection. Such correlations might reflect differences in the host immune response dynamics or treatment-related modulation of the disease course. Further investigations using longitudinal data and viral load measurements would help elucidate these differences.

**Fig 4.**
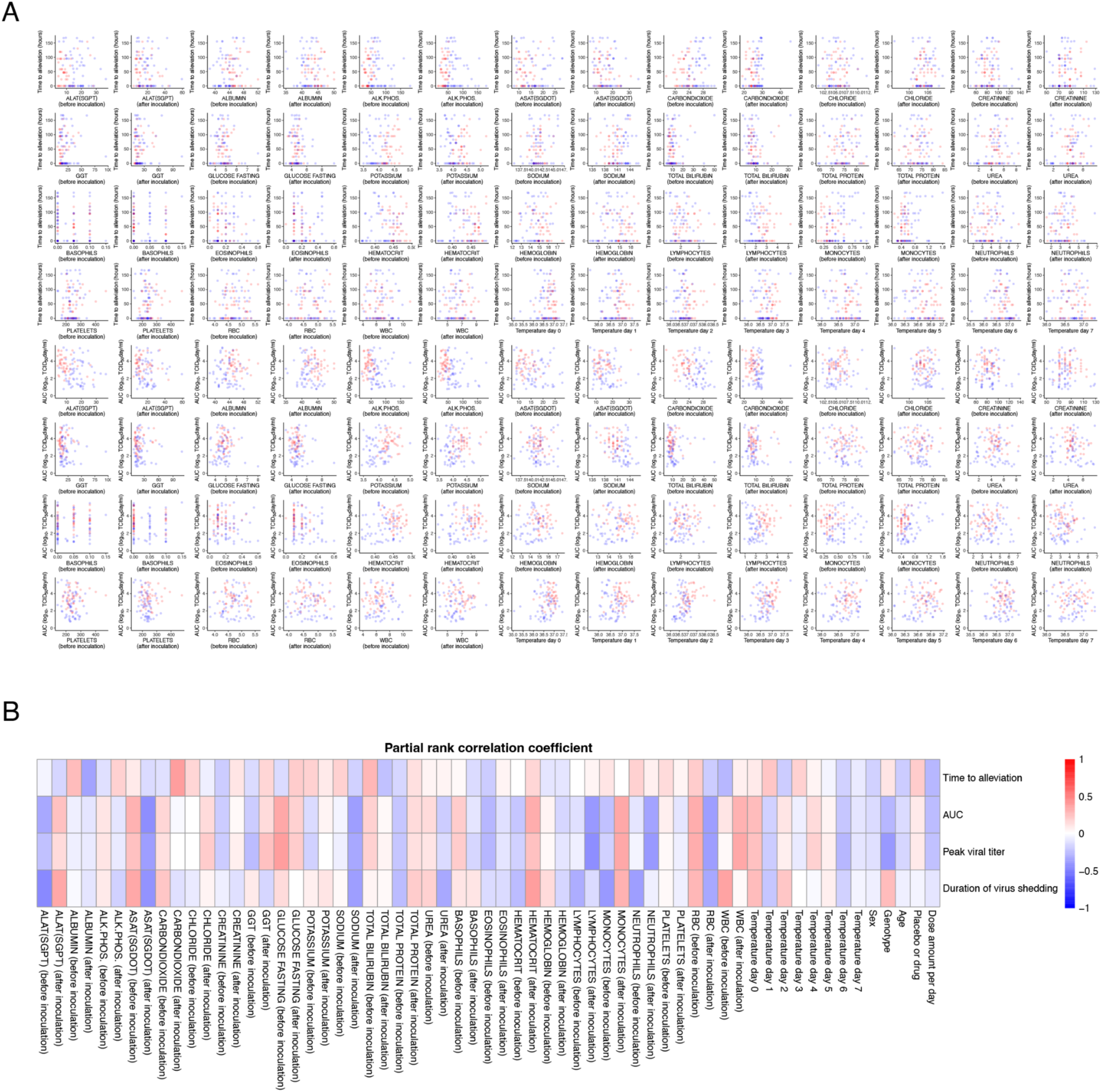
Correlations between participant status and outcomes. **(A)** Relationships between the time to alleviate composite symptoms and laboratory measurements and body temperature (upper half) and between the AUC and laboratory measurements (lower half). **(B)** Partial rank correlation coefficients of participant background and clinical measurements against time to alleviation, AUC, peak virus titer, and duration of infection.

## Discussion

Using mathematical modeling and statistical approaches, we analyzed several data types extracted from three clinical trials of oseltamivir treatment of experimental influenza infection. Although oseltamivir has been shown to suppress symptoms and viral shedding in clinical trial outcomes, our findings indicate the possibility of unclear and different effects of oseltamivir administration on the outcomes.

Using mathematical models of viral infection dynamics to analyze the viral load allowed for more interpretable comparisons of antiviral efficacy. Our primary findings indicated that oseltamivir-treated individuals tended toward lower viral load AUC, shorter duration of infection, lower peak viral titers, and earlier time to peak than those receiving a placebo. These changes may be caused by accelerated viral infection and replication. However, most of these differences were not significantly different, and no apparent dose-dependent effects were observed. The significant advantage of high-dose efficacy has not been confirmed in multiple clinical trials or meta-analyses [36, 42–44].

Notably, no significant correlation was found between viral load and time to symptom alleviation. The weak negative correlations of laboratory test data with symptoms and viral load suggested a different mechanism of action for oseltamivir, but no apparent causal relationship was obtained. Oseltamivir, known as a neuraminidase inhibitor, has been shown through experimental observations to alleviate symptoms not by inhibiting neuraminidase but through other mechanisms [4, 16, 45], and it is considered likely that the same applies in humans. Notably, laboratory tests were performed in the trials to assess the safety of oseltamivir treatment and were not expected to change with or without oseltamivir administration. The three trials from which data were obtained and others reported no change in laboratory safety evaluations at the population level except for a small subset of participants [35, 36, 40, 41].

Unlike previous meta-analyses based on summary statistics, our present study reanalyzed individual-level viral load and laboratory test measurements from published and available data from clinical trials. A single endpoint may not fully capture the therapeutic effect of oseltamivir because some participants recovered from symptoms without suppressed viral shedding. In addition, individual symptom transitions remain a subject of investigation because time to symptom alleviation obscures differences in symptom combinations among participants. Furthermore, no data on the immune response were obtained, and viral elimination or inflammatory response could not be assessed.

For pandemic preparedness and response, public health aspects of treatment, such as the decreased chance of infection due to sustained viral shedding, need to be evaluated, in addition to the impact on individual symptoms. In clinical use and stockpiling for a pandemic, the usefulness of antiviral drugs must be considered in the evaluation. Our results highlight the need for more refined clinical trial endpoints, enhanced public access to individual patient data, and further investigations into the impact of oseltamivir on viral transmission. Future studies must explore mechanism-of-action-based dosing strategies, alternative antiviral strategies, and combination therapies to achieve more optimized influenza treatment.

## Limitations

Notably, the present study was not intended to evaluate oseltamivir drug efficacy; instead, it was an exploratory analysis. In particular, we excluded data from participants from the influenza-infected population who did not have sufficient viral shedding to estimate viral transmission dynamics. The estimated parameters of the mathematical model also need to be carefully considered to interpret the possible increase in viral infection and viral replication as an effect of oseltamivir administration. Because of the nature of our target-limited model, the faster depletion of target cells by promoted infections may have led to a quicker decrease in infection and may explain the reduction in viral shedding.

## Methods

### Clinical trials for which datasets were obtained

We used datasets from two randomized controlled trials to assess the efficacy of oseltamivir treatment against experimental influenza virus A and B infection [40, 41]. For the influenza A study (Flu A study), the 80 participants were randomized to receive one of five treatments (20 mg b.i.d., 100 mg b.i.d., 200 mg b.i.d. or 200 mg o.d. oseltamivir, or placebo). Participants were inoculated with 10^6^ median tissue culture infectious doses (TCID_50_) of human influenza virus A/Texas/91 (H1N1) and administered oseltamivir or placebo for 5 days beginning 28 h after virus inoculation. In study A on the trial against influenza B infection (Flu B study A), the 60 participants were randomized to receive one of three treatments (75 mg b.i.d. or 150 mg b.i.d. oseltamivir or placebo). Participants were inoculated with 10^7^ TCID_50_ of human influenza B/Yamagata/16/88 virus. Oseltamivir or placebo was administered 24 h after viral infection for 5 days. In study B of the trial against influenza B virus (Flu B study B), the 117 participants were randomized to receive one of two treatments (75 mg b.i.d. oseltamivir or placebo). The inoculation of influenza virus followed the same protocol as Flu B study A. We included data from 69, 39, and 86 participants who received treatment in the efficacy analysis of oseltamivir treatment in the Flu A study, Flu B study A, and Flu B study B, respectively (**Table S7**). The primary endpoint was the change in the AUC of virus titer in both the Flu A and Flu B studies. Peak viral load or duration of viral shedding from treatment initiation were evaluated as secondary endpoints related to the viral load. In addition, symptom scores and time to alleviate total or composite symptoms were compared with symptom-based assessments in natural infection [35, 36].

### Digitalization of measurements in clinical trials

To assess the effect of oseltamivir treatment on influenza virus infection, background information on the participants (i.e., age and sex), items measured in clinical trials (i.e., standard clinical laboratory tests and temperature), and time to alleviation of symptoms were obtained as numerical data from PDF files of protocol numbers PV15616 for the Flu A study, NP15717 for the Flu B study A, and NP15827 for the Flu B study B, on a published database [6]. Because the definitions of time-based virus inoculation differed between the Flu A and Flu B studies, the time of inoculation was defined as the first half day of day 0 in our present study (**Figure S1**). In detail, in the Flu A study, participants were inoculated with viruses on the afternoon of day 1, and the data were collected from the morning of day 2 to the morning of day 9 with the date defined in the trial. In the Flu B study, the date of inoculation was the morning of day −1, and data were collected from the afternoon of day −1 to the morning of day 8. Day 0 did not appear in the timeline. To facilitate the combined analysis of data from both studies, we defined the time at 24 h following inoculation as day 0.

Background information on the participants is listed in **Table S7**. Laboratory test data were obtained from hematologic and biochemical measurements performed in the Flu A and B studies. Urinalysis data were excluded from the analysis in the present study because there were insufficient data in the Flu B study. The times and measurements of the laboratory tests for individual participants are summarized in **Tables S1–S3**. Body temperatures were measured 4 times a day and 3 times for preliminary and follow-up testing, 32 times in the Flu A study and twice a day and 3 times for preliminary and follow-up testing, 20 times in the Flu B study (**Tables S4–S6**).

The time to alleviate seven influenza symptoms, cough, nasal obstruction, sore throat, fatigue, headache, myalgia, and feverishness, was defined as the first time from the start of treatment to the subsequent time at which all the symptoms had reduced to absent or mild. Participants with at least one or more symptoms that were not mild at the final assessment were censored. We digitized the time to alleviate seven symptoms reported for each participant from the PDF files and used them for our analysis (**Table S7**).

### Viral load data

The viral load data from the oseltamivir trials were digitized and published previously [39]. The published dataset included the viral load with a summary of information about participants and procedures for each study. For the parameter estimation in the mathematical model of viral infection dynamics, we excluded data from 91 of the 194 participants included in the efficacy analysis in the clinical trials with less than 3 points of viral load data above the detection limit (**Figure 1A** and **Table S7**). Notably, all participants excluded from the efficacy analysis in the clinical trials met the above exclusion criteria defined by the number of viral load observations.

### Aggregation of measurements for correlation analysis

The items determined in the laboratory tests were hemoglobin (g/dL), hematocrit (fraction), RBC (10^12^/L), WBC (10^9^/L), neutrophils (10^9^/L), lymphocytes (10^9^/L), monocytes (10^9^/L), eosinophils (10^9^/L), basophils (10^9^/L), platelet count (10^9^/L), sodium (mmol/L), potassium (mmol/L), chloride (mmol/L), urea (mmol/L), creatinine (μmol/L), total protein (g/L), albumin (g/L), blood glucose (mmol/L), phosphate (mmol/L), uric acid (μmol/L), triglycerides (mmol/L), cholesterol (mmol/L), AST (SGOT) (U/L), ALT (SGPT) (U/L), GGT (U/L), alkaline phosphatase (U/L), total bilirubin (μmol/L), calcium (mmol/L), LDH (U/L), carbon dioxide (mmol/L), CPK (U/L) and serum amylase (U/L) for Flu A study and hemoglobin (g/dL), hematocrit (fraction), RBC (10^12^/L), WBC (10^9^/L), neutrophils (10^9^/L), lymphocytes (10^9^/L), monocytes (10^9^/L), eosinophils (10^9^/L), basophils (10^9^/L), platelet count (10^9^/L), MCHC (g/L), MCV (fL), sodium (mmol/L), potassium (mmol/L), chloride (mmol/L), urea (mmol/L), creatinine (μmol/L), total protein (g/L), albumin (g/L), AST (SGOT) (U/L), ALT (SGPT) (U/L), GGT (U/L), alkaline phosphatase (U/L), total bilirubin (μmol/L), blood glucose (mmol/L), and carbon dioxide (mmol/L) for Flu B study (**Tables S1–S6**). We selected 24 items of the measured items every day for the Flu A and Flu B studies for subsequent analysis (**Table S7**). These laboratory measurements had been made 2 times before virus inoculation and once at 8 days after virus inoculation in the Flu A study and once before virus inoculation and twice at 3 days and 5 days after virus inoculation in the Flu B study (**Tables S1–S6** and **Figure S1**). The data from the laboratory tests were split before and after virus inoculation to ensure consistent analysis in the Flu A and Flu B studies. The means of the values were calculated if two measurements were performed (**Table S7**). The average daily body temperatures (degrees Celsius) after virus inoculation were calculated for correlation analysis (**Table S7**). Body temperature on day 8 after virus inoculation was excluded because it was not measured in the Flu A study. Among the 103 participants who met the viral load analysis criteria, data from 2 who were missing one or more items either before or after virus inoculation were excluded from the plots in **Figure 1** and from the correlation analysis (**Figure S1** and **Table S7**).

### Mathematical model of virus dynamics

To analyze the viral load of the influenza virus in the participants, the following mathematical model of within-host virus infection dynamics with quasi-steady state approximation [46–50] was employed:

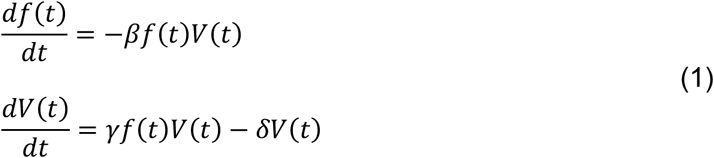

The variable *f*(*t*) represents the fraction of target cells at time *t* relative to the baseline value at time 0, and the variable *V*(*t*) represents the number of viruses at time *t*. The parameters *β* and *γ* represent the scaled rate constant for infection and the maximum rate constant for viral replication, respectively. Viruses are eliminated, and the death rate of infected cells is *δ*.

### Parameter estimation of a mathematical model

Unknown parameters, *β*, *γ*, *δ*, and *V*(0), in the model were estimated from the viral load data. Because the virus was not detected in all participants on day 0 after virus inoculation and the intervention was performed from day 1, data from day 0 were excluded from the data fitting. The model parameters were estimated using a nonlinear mixed-effects model implemented in Monolix [51] to account for variation in viral load changes across participants. The parameters for each participant were assumed to follow a log-normal distribution, and genotypes were used as covariates. The values of the population parameters estimated using a stochastic approximation expectation-maximization algorithm and the individual parameters estimated as a mode of the conditional parameter distribution calculated as empirical Bayes estimates are listed in **Tables S8** and **S9**, respectively.

### Viral load-related outcomes estimated by a mathematical model

The AUC, duration of virus shedding, peak viral titer, and time to peak viral titer were calculated using viral load estimated by a mathematical model from treatment initiation, i.e., 1 day after virus inoculation, until the viral load dropped below the detection limit. In accordance with a previous study, the detection limit was set as 1 TCID_50_/mL [39].

### Partial rank correlation coefficients (PRCCs) between outcomes and participant status

The PRCCs of the laboratory test data, sex, age, infected genotype, treatment arms, and endpoint dose were calculated using the R package ppcor [52].

## Data availability

Viral load data are publicly available from a publication [39]. Other data are available within this article as **Supplementary Tables**.

## Code availability

Zenodo provides codes for estimating model parameters with a nonlinear mixed-effects model [53].

## Acknowledgments

We thank Robin James Storer, PhD, from Edanz (https://jp.edanz.com/ac) for editing a draft of this manuscript.

## Author contributions

**Conceptualization**, Y.F., N.N., and S. Iwanami; **Methodology**, S. Iwami, N.N., and S. Iwanami; **Investigation**, Y.F., M.A., D.T., N.N., and S. Iwanami; **Data Curation**, Y.F., M.A., and D.T.; **Writing – Original Draft**, Y.F., M.A., D.T., N.N. and S. Iwanami; **Writing – Review & Editing**, S. Iwami, N.N., S. Iwanami; **Visualization**, Y.F., M.A., D.T., and S. Iwanami; **Supervision**, S. Iwami, N.N., and S. Iwanami; **Funding Acquisition**, S. Iwami, and S. Iwanami.

## Supplemental information

### Supplementary Figures

**Fig S1.**
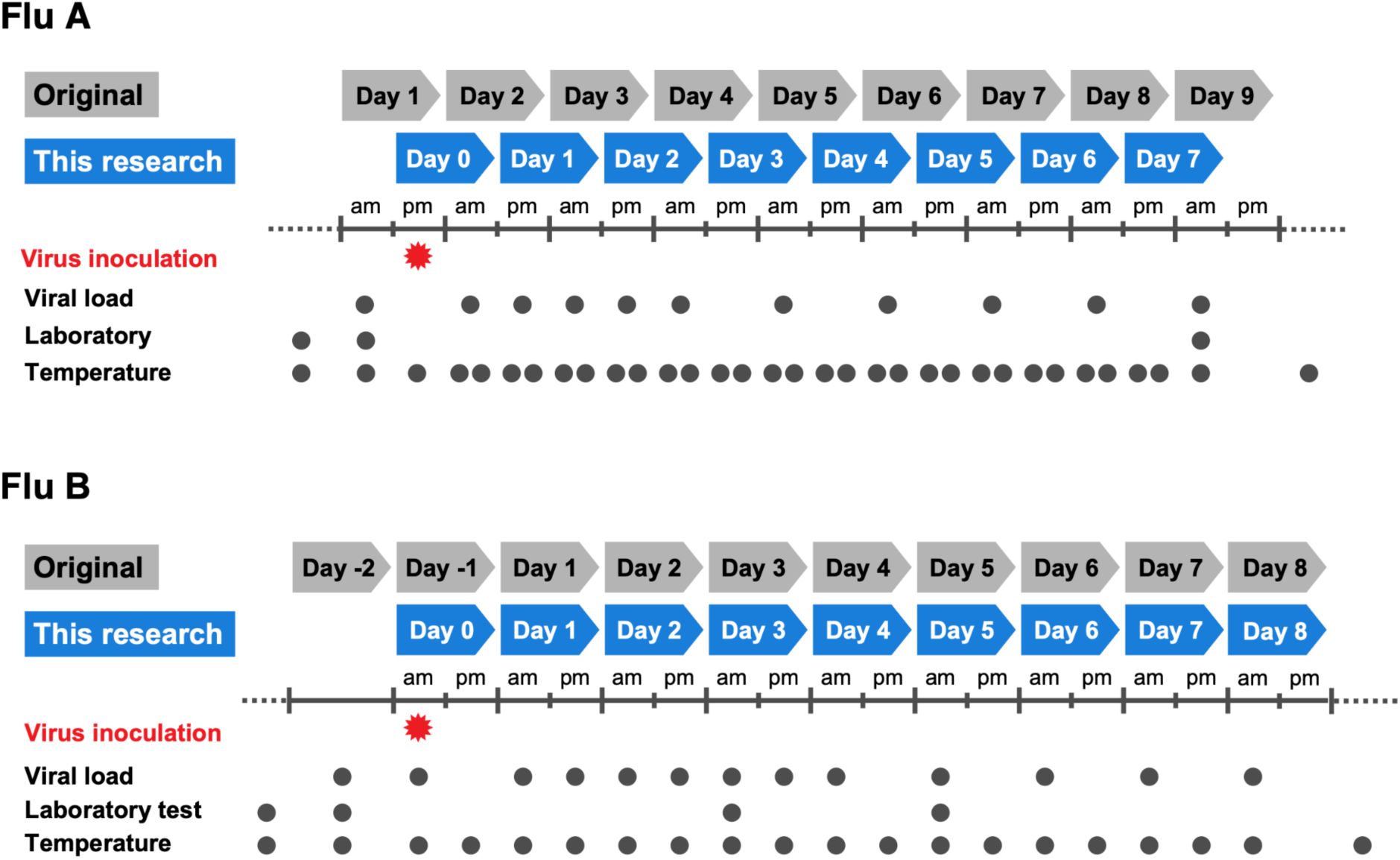
Measurement schedule of data in clinical trials. The schedule from which each datum point used in the analysis was measured in the clinical trials. Times were modified according to the time of inoculation and used for the data analysis.

**Fig S2.**
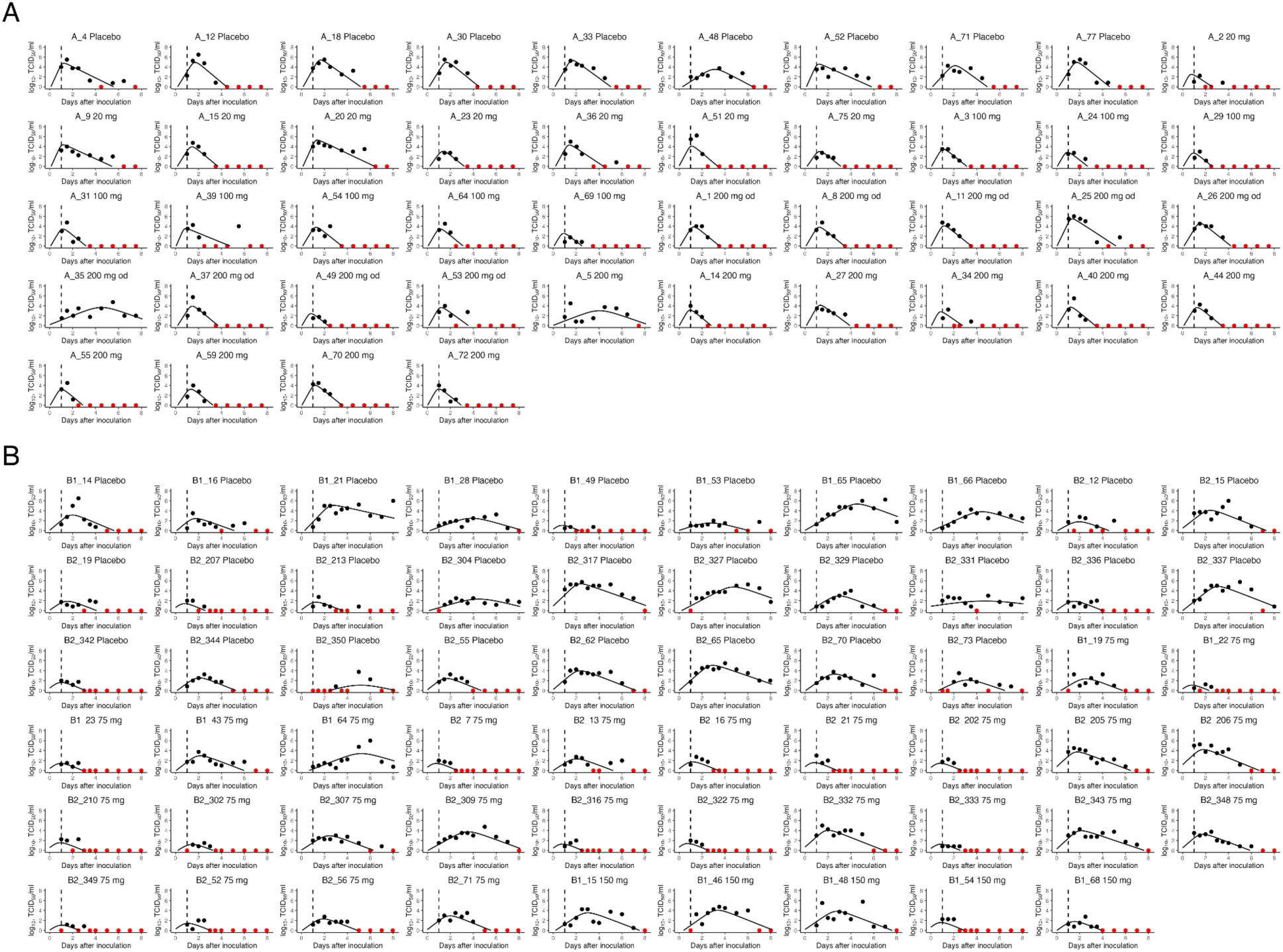
Individual model fit to viral load data. (A and. **B)** Measured viral load data (dots) and viral load calculated by a mathematical model using parameters estimated by a nonlinear mixed-effect model (lines) for influenza A **(A)** and B **(B)**, respectively. The red dots represent viral titers under the detection limit. The doses without “od” were administered twice daily; the dose with “od” was administered once daily.

**Fig S3.**
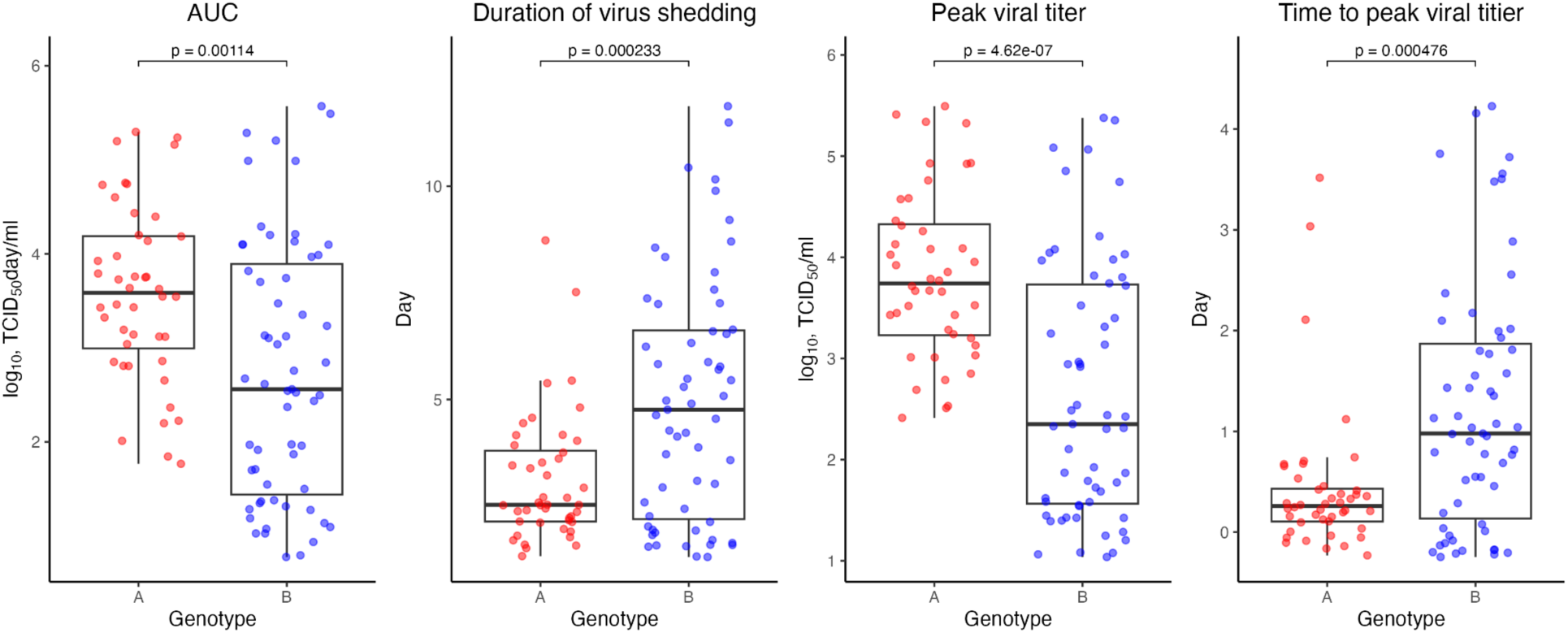
Comparison of estimated viral load features between influenza A and B virus infections. Comparison of estimated viral load-related outcomes, AUC, duration of virus shedding, peak viral titer, and time to peak viral titer between influenza A and B infection. The means were compared using a *t* test, and *p*-values were adjusted using a post hoc Bonferroni correction.

**Fig S4.**
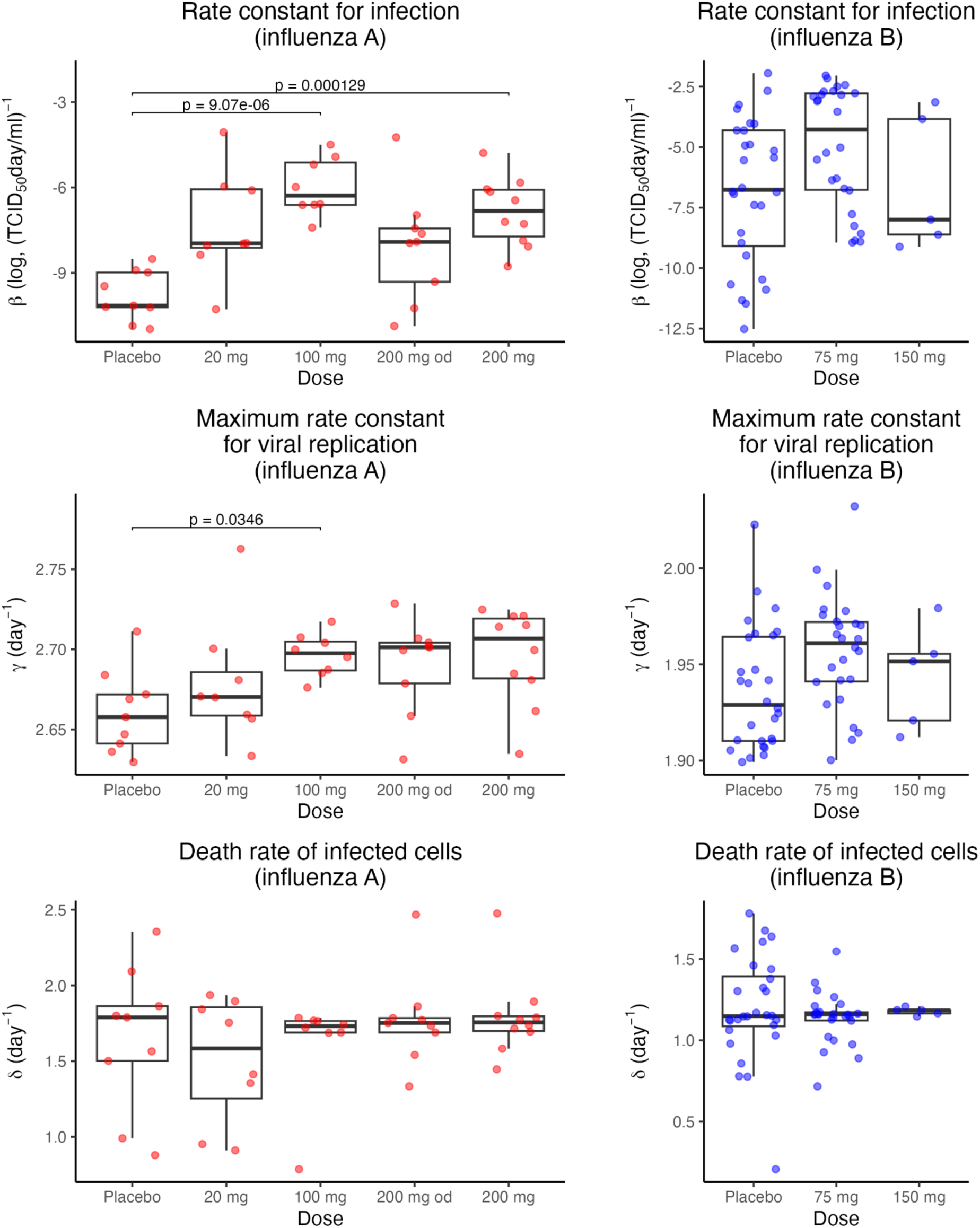
Relationships between estimated parameters and viral strains and oseltamivir treatment. Comparison of the estimated model parameters between groups by dose of oseltamivir for influenza A (left) and influenza B (right) viruses. The doses without “od” were administered twice daily; the dose with “od” was administered once daily. We adjusted *p*-values according to the number of group combinations using a post hoc Bonferroni correction.

### Supplementary Tables

**Table S1. Laboratory test data from the Flu A study.**

Original measurements for blood chemistry and hematology extracted from standard laboratory test results in the Flu A study are listed in **TableS1.csv**.

**Table S2. Laboratory test data from the Flu B Study A.**

Original measurements for blood chemistry and hematology extracted from standard laboratory test results in the Flu B study A are listed in **TableS2.csv**.

**Table S3. Laboratory test data from the Flu B Study B.**

Original measurements for blood chemistry and hematology extracted from standard laboratory test results in the Flu B study B are listed in **TableS3.csv**.

**Table S4. Body temperature in the Flu A study**

Original body temperature for each participant extracted from the Flu A study is listed in **TableS4.csv.**

**Table S5. Body temperature in the Flu B Study A**

Original body temperature for each participant extracted from the Flu B study A is listed in **TableS5.csv.**

**Table S6. Body temperature in the Flu B Study B**

Original body temperature for each participant extracted from Flu B study B is listed in **TableS6.csv.**

**Table S7. Participant information, symptoms, laboratory data, and time courses of temperature in the human challenge**

The participant information, i.e., ID, age, sex, study, genotype of the inoculated virus, treatment, exclusions in each analysis, and extracted values of time to alleviation, laboratory data, and temperature, are listed in **TableS7.csv**.

**Table S8.** Model parameters estimated by a nonlinear mixed-effects model.

| Name | Symbol | Unit | Population value | Variance of individual parameters | Covariate for genotype B (p-value by Wald test) |
| --- | --- | --- | --- | --- | --- |
| Rate constant for infection | $\beta$ | (RNA copies/mL)<sup>-1</sup>day<sup>-1</sup> | $5.55 \times 10^{-4}$ | 2.68 | 1.65<br>($1.37 \times 10^{-2}$) |
| Maximum rate constant for viral replication | $\gamma$ | Day<sup>-1</sup> | 14.7 | $9.55 \times 10^{-2}$ | -0.739<br>($2.47 \times 10^{-13}$) |
| Death rate of infected cells | $\delta$ | Day<sup>-1</sup> | 5.18 | 0.411 | -0.595<br>($2.98 \times 10^{-3}$) |
| Initial value of viral load | V(0) | TCID<sub>50</sub>/mL | 0.492 | 1.96 | 0.959<br>(0.659) |

**Table S9. Estimated individual parameters and features related to viral load**

Values of estimated individual parameters, *β*, *γ*, *δ*, *V*(0), features related to viral load, area under the curve, duration of infection, peak viral titer, time to peak viral titer, and daily viral load, calculated with estimated individual parameters are listed in **TableS9.csv**.

